# Divergence in wilt tolerance across the range of *Mimulus guttatus* in the absence of stomatal regulation

**DOI:** 10.64898/2026.09.30.755782

**Authors:** Haley A. Branch, Pauline Raimondeau, Vanessa Tonet, Henry Arenas Castro, Andrew Fairclough, Lily Hyde, Dristen Lakes, Alyssa Singletary, Benjamin K. Blackman, Katherine Toll, Mario Vallejo-Marín, Craig Brodersen, Erika Edwards, Jenn M. Coughlan

## Abstract

Whether phenotypes and environments predict performance under stress is central to ecology and evolution and can reveal species’ vulnerability to global change. We sampled *Mimulus guttatus* range-wide to test whether reproductive, morphological, and physiological traits predict drought responses. Surprisingly, we find that stomatal response, a key mediator of water balance, did not change as drought intensified. Climate, however, predicted survival after turgor loss: plants from regions with large daily relative to seasonal temperature ranges survived wilting longer. Stress responses also depended on life history: where summers are dry, perennials withstood lower water potentials, whereas annuals increased leaf cooling as they wilted. Known trait associations were not fixed: although many baseline traits tracked climate, none predicted drought responses. Drought performance therefore cannot be inferred from traits measured under well-watered conditions. Lacking stomatal regulation, riparian drought escapers like *M. guttatus* may be vulnerable if shorter growing seasons bring drought before reproduction.

## Introduction

Plant ecologists have long sought to characterize functional traits and how they relate to underlying physiological mechanisms (Farrell et al., 2017; Funk et al., 2024; Branch, 2026). Such trait-trait associations allow predictive modeling of species’ responses to their environment and have improved our understanding of global patterns in plant form and function (Díaz et al., 2016). Evolutionary biologists have long examined these associations as trade-offs and constraints in local adaptation, using them to glean how species might rapidly evolve with climatic change (Griffiths et al., 2024). While patterns such as the leaf economic spectrum have been found to hold across taxa (Anderegg et al., 2018), evidence suggests these relationships can be more complex and once-conserved associations can be uncoupled (Edwards et al., 2014; Anderegg et al., 2018; Kooyers et al., 2015; Schwab et al., 2026). This poses a significant issue for understanding how plant populations will respond to the drier, more variable environments of the future.

Plant evolutionary responses to drought reveal common trait–trait associations, represented as functional groups mediated by life history and water conservatism. Herbaceous plants are often grouped into three drought-response strategies: escape, avoidance, and tolerance (Chaves et al., 2003; Ludlow & Muchow, 1990; Kooyers, 2015; Anstett et al., 2021; Branch et al., 2024a; Branch et al., 2024b; Schwab et al., 2026). Escape involves rapid development and reproduction before drought onset and is associated with an annual life history and risky water use via high stomatal conductance and growth rates (Levitt, 1980; Chaves et al., 2003; Franks, 2011; Kooyers, 2015). These plants are highly resource-acquisitive, with larger, cheaper leaves and little water conservatism (Gonzalez-Paleo & Ravetta, 2018; Funk et al., 2024). Avoidance involves slower growth, reproduction at later nodes, reduced stomatal conductance, and behavioural (e.g., leaf rolling and shedding) and morphological changes (e.g., stomatal size, succulence; Chaves et al., 2003; Kooyers, 2015; Kramp et al., 2022). Avoidant plants are highly water-conservative, rapidly and fully closing stomata to maintain whole-plant turgor (Bartlett et al., 2016), which is effective only as long as they can forgo carbon assimilation (Guadagno et al., 2017). Tolerance relies on biochemical and structural traits that limit drought damage, such as high capacitance (water storage), osmotic adjustment, or resistant xylem, allowing plants to sustain riskier, escape-like water-use, growth and reproduction (Chaves et al., 2003; Kooyers, 2015; Bartlett et al., 2016; Guadagno et al., 2017; Nolan et al., 2017; Kramp et al., 2022). Perennials are generally avoidant or tolerant and often trade-off water-use efficiency against growth (Zeri et al., 2013; Valliere, 2019). Their leaves are thicker and costly and may be more dissected, increasing convective cooling and water-use efficiency (Chaves et al., 2003; Vogel, 2009; Nicotra et al., 2011; Kooyers, 2015; but see Schwab et al., 2026). These strategies are not mutually exclusive within species, and widespread species likely vary in strategy across their ranges (Anstett et al., 2021; Funk et al., 2024; Branch et al., 2024b; Schwab et al., 2026). Thus, wide-ranging species, and especially those with both annual and perennial life histories, are excellent candidates for examining differences in water-conservation strategies and drought responses.

An often-overlooked area of vulnerability to climate change are riparian plants that rely on ephemeral rivers fed by precipitation and snowmelt. With roots generally in direct contact with water, these species are characterized by a drought-escape strategy. However, projected changes in precipitation patterns and reduced snowpack will shorten the growing season for downstream species (Zimmer et al., 2022). Riparian growing seasons generally end with a terminal drought (see Box 1) after reproduction, inducing senescence or dormancy. Such species likely lack drought avoidance or tolerance adaptations, unlike species that experience drought throughout their growing season. Understanding riparian plant responses to rapid-onset terminal drought will shed light on their vulnerability to future environments.

Here we combine range-wide variation, divergent life histories, and riparian vulnerability, using extensive sampling of perennial and annual *Mimulus guttatus* (syn. *Erythranthe guttata*) accessions to assess phenotypic differences and drought response in this monkeyflower. We predicted that northern populations, where water availability is more continuous (Fig. S1), would exhibit escape, while southern populations, where water availability is lower, would exhibit avoidance or tolerance. We also predicted that perennials would have greater capacity to avoid or tolerate drought than annuals, which would be more likely to escape. Our results shed light on the vulnerability of riparian species and provide nuance to the leaf economic spectrum.

## Methods

### Study system, collection, and growth

*Mimulus guttatus* DC (syn. *Erythranthe guttata* Nesom 2014; Barker et al., 2012; see Lowry et al., 2019; Nesom et al., 2019) is a C3 herbaceous riparian plant with annual and perennial life histories (Hall & Willis, 2007; Lowry & Willis, 2010; Friedman et al., 2015). It ranges across western North America, from Alaska’s Aleutian Islands to Mexico (Coughlan et al., 2021). We obtained 658 accessions from 306 populations spanning the species’ range (Fig. 1).

**Figure 1.**
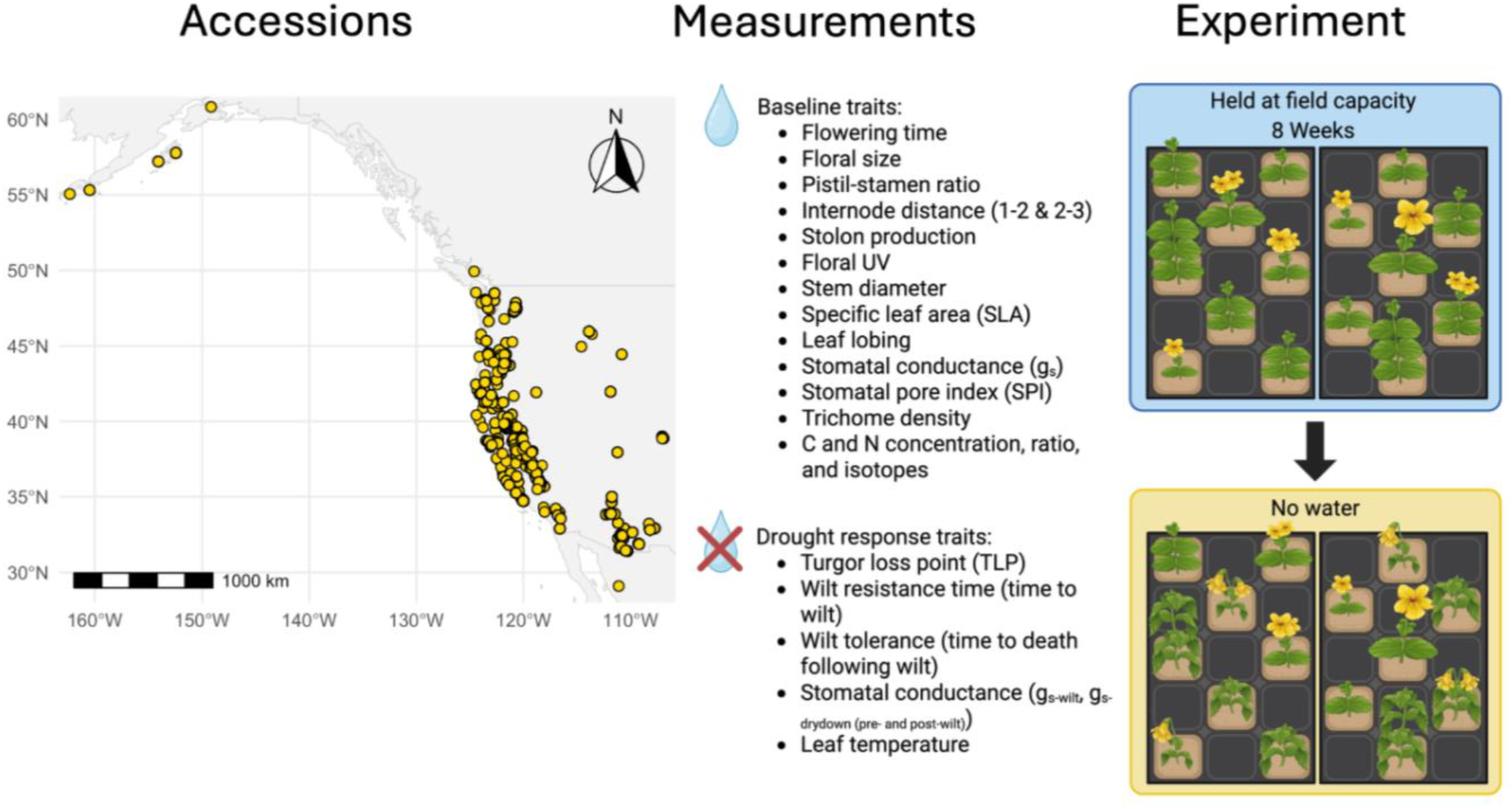
Accession collection and experimental design. We grew 658 accessions of *Mimulus guttatus* from across its range in a common greenhouse. For eight weeks, plants were held at water saturation and baseline traits were measured. After eight weeks, we implemented a dry-down to simulate terminal drought, which is characteristic of ephemeral rivers and measured drought response until senescence.

Field-collected annual and perennial populations were refreshed for one generation to reduce parental effects. Seeds were cold-stratified in distilled water in Eppendorf tubes at 4℃ for 5 days, then sown in 2.5” pots of Promix soil (same sowing conditions throughout unless stated) in the Yale plant growth facilities greenhouse. The greenhouse was set to 25℃ with 16-hr days, and benches were flooded daily to saturate the soil. At reproductive maturity, we performed one round of random crosses within populations, emasculating buds the day before opening to prevent autogamy. Populations with only one or two accessions were refreshed by selfing.

Seeds from refreshed accessions were stratified as above, then germinated in growth chambers at 21℃ (day and night) with 16-hr days. Single seedlings were transplanted into fresh 4” pots and placed in one of two greenhouses (∼22℃ day and night, 16-hr days, natural light supplemented with incandescent lighting as needed) in a randomized block design. For their first eight weeks, plants were bottom-watered several times daily to keep the soil saturated; watering then ceased to begin the dry-down (Fig. 1). By growing one seedling per pot at constant pot and soil volume, we aimed to identify range-wide phenotypic differences attributable to genetic differences. However, germination time varied (difference of 0-21 days, mean 1.58 days) and plants entered the dry-down at different developmental stages and sizes.

### Measurements

#### Reproductive traits

We recorded the date and node of the first flower. The first flower was removed on the day it opened to measure flower length, corolla width, corolla tube length, longest stamen length, and pistil length. Selfing potential was measured as the distance between the longest stamen and the pistil. Stolons were also counted at this time to ensure the same developmental stage across accessions. On another flower, we measured floral UV absorption with a Canon EOS Rebel T7 DSLR camera fitted with a custom UV band-pass filter in place of the hot-mirror (LifePixel, WA). Using ImageJ v. 1.53a (Schneider et al., 2012), we quantified UV absorption as the proportion of the interior medial ventral lobe and corolla tube that was UV-absorbent (i.e., absorbent length/total floral length). We examined UV absorption because it varies with latitude and elevation within species (Koski & Ashman, 2016; Norbona et al., 2025), putatively differs between annual and perennial *M. guttatus* (DeMarche et al., 2015), and has a cost that could affect whole-plant resource use.

#### Morphological and physiological traits

On the day the first flower opened, we also measured stem diameter and internode length between the first and second nodes (growth), and internode length between the second and third nodes (growth and bolting). Baseline stomatal conductance (gs-baseline) was measured on one third-node leaf between 0800 and 1200 using a LI-600 porometer (LI-COR Inc., Lincoln, NE, USA). This leaf was then removed to measure specific leaf area (SLA = one-sided leaf area/leaf dry mass). Leaf area was measured in ImageJ from scanned images (Blonder, accessed 2024). Leaves were oven-dried at 60℃ and weighed. The same leaves were then ground with a Spex 2010 Geno/Grinder, and carbon and nitrogen isotope ratios were measured with a Thermo DeltaQ isotope ratio mass spectrometer coupled to an EA Isolink CN. Leaf margins in the scanned images were processed in ImageJ and Python to determine percent lobing (Lamb, 2025).

The second third-node leaf was used for epidermal peels. Trichomes were first removed gently with labeling tape, then a thin layer of clear nail polish was applied to the abaxial surface (although *Mimulus* also has adaxial stomata) between the midvein and margin at the leaf center. Once the polish had cured, peels were imaged under a compound light microscope. Removing trichomes first keeps them off the stomata, allows the polish to cure, and leaves a circular mark at each trichome base that is distinguishable from stomata and countable. Trichome and stomatal densities (per mm^2^) and guard cell length were measured in ImageJ. Stomatal pore index was calculated as SPI = SD × SL2, where SD is stomatal density and SL is stomatal length (Liu et al., 2023).

#### Drought traits

Before the dry-down, a fourth-node leaf was collected between 0700 and 1100 hr to measure turgor loss point (Ψ_TLP_). A 5 mm leaf disc was wrapped in aluminum foil, flash-frozen in liquid nitrogen, and pierced with fine forceps. The remaining leaf tissue was weighed fresh and dry to obtain leaf dry matter content (LDMC). Osmotic potential was measured with a vapor pressure osmometer (VAPRO 5600, EliTech Inc., Logan, UT, USA) and used with LDMC to calculate Ψ_TLP_ (Bartlett et al., 2012; Petruzzellis et al., 2019).

During the dry-down, plants were checked for wilting and senescence at 0900 and 1600 hr. Plants were scored as wilted when leaves lost turgor and as dead when they became desiccated and necrotic. This yielded wilt resistance (days from drought initiation to wilting) and wilt tolerance (days from wilting to death). In a subset of 222 plants, stomatal conductance and leaf temperature were measured with the LI-600 porometer to examine stomatal regulation and thermoregulation. Measurements were taken at 0800–1000 and 1300–1500 on ten consecutive days, until all plants had died, to capture gs variation within days and across the dry-down. Because each session spanned two hours, air temperature and solar radiation (not measured) likely varied, particularly in the morning. Stomatal conductance at wilting was recorded to relate conductance to wilt resistance and tolerance.

### Statistical analyses

#### Climate models

We built a maximum entropy species distribution model in MaxEnt v3.4.4 (Phillips et al., 2006; Da Re et al., 2020) from our sampling sites and 3,189 *M. guttatus* occurrence records from the Global Biodiversity Information Facility (GBIF; https://doi.org/10.15468/dl.nkfajq), restricted to the native range (longitude −180° to −60°). We evaluated the relative importance of 19 bioclimatic variables from WorldClim v2.1 (Fick & Hijmans, 2017) at 2.5 arc-minute (∼4.5 km) resolution, assessing performance by five-fold cross-validation and the area under the receiver operating characteristic curve (AUC) of each replicate. A PCA (*prcomp*) of our accessions’ WorldClim data identified the variables contributing most variation, and together the two approaches yielded four variables (Table S1; Fig. S2): isothermality (bio3), temperature seasonality (bio4), annual precipitation (bio12), and precipitation of the warmest quarter (bio18).

#### Data curation

Of the 28 measured traits, we retained one of each pair correlated above 0.65 (*rcorr*, ‘Hmisc’; Harrell & Dupont, 2019; R v4.5.2, R Core Team, 2025; Fig. S3). Five floral size traits (total flower length, corolla width, corolla tube length, stamen length, and pistil length) were reduced to one principal component (88.1% of variance; *prcomp*). Of the remaining 18 traits, we removed one of each pair with a variance inflation factor (VIF) > 10, which excluded only the carbon-to-nitrogen ratio (VIF > 20). The same procedure reduced five drought traits to four, as wilt resistance and senescence were highly correlated.

#### Trait-trait associations across the range

For each of the four drought responses, we fit a least absolute shrinkage and selection operator (LASSO) regression (‘glmnet’; Friedman et al., 2010) with the 17 remaining traits as predictors. Missing predictor values (26% for stomatal pore index, 25% for leaf lobing, <20% for all others) were imputed by predictive mean matching (‘mice’; van Buuren & Groothuis-Oudshoorn, 2011; Table S2); rows missing the response were dropped rather than imputed. For each response, we compared raw, log, and square-root scales using Q–Q plots and histograms of residuals, Shapiro– Wilk tests, and skewness, and used the least-skewed scale (Table S3).

The number of imputations was set with *how_many_imputations* (von Hippel, 2020), targeting a coefficient of variation of 0.05 (reproducible without being overly strict) and a significance threshold of 0.09 (more flexible given the number of predictors). Variables were standardized internally and batch (greenhouse 1 or 2) was an unpenalized fixed effect (*penalty.factor*). To assess selection stability, we drew 200 bootstrap samples from each imputed dataset, chose each resample’s penalty by 10-fold cross-validation (lambda.1se), and used a selection-frequency cut-off of 0.60. We ran the same LASSO with gs-baseline as the response and the other 16 traits as predictors. Selected traits were then tested in linear mixed models (‘lme4’; Bates et al., 2015) with batch as a fixed effect and population, greenhouse bench, and pot position nested within tray as random effects, using Type III ANOVA (‘car’; Fox & Weisberg, 2019) and Benjamini–Hochberg (BH)-corrected *p*-values.

#### Stomatal conductance during wilt

We used repeated-measures mixed models, indexing each measurement by time point (*x*), to ask three questions about stomatal conductance, thermoregulation, and their climate associations during the dry-down. (1) Does climate predict the g_s-drydown_ trajectory before and after wilting? Separate pre- and post-wilt models used g_s-drydown_ at *x* as the response, with wilt resistance or wilt tolerance (days), respectively, interacting with each of the four biovars, which did not interact with one another. (2) Does the coupling of leaf temperature and g_s-drydown_ (thermoregulation) differ before and after wilting? Here, leaf temperature at *x* was modelled against within-plant g_s-drydown_ at *x* and its interaction with wilt stage (pre- or post-wilt), with mean gs and relative time point as covariates. (3) Does climate predict the leaf temperature–g_s-drydown_ relationship within wilt phase? Separate pre- and post-wilt models related leaf temperature at *x* to interactions between within-plant g_s-prewilt_ or g_s-postwilt_ and each of the four biovars, testing whether plants stop thermoregulating before wilting and whether they transpire passively after it. All models used batch and random-effect structure above, Type III ANOVA (‘car’), and BH-corrected *p*-values.

#### Climate-trait associations

To test climate associations, we fit 21 additive linear mixed models, one per baseline trait and drought response, with the four z-scored biovars (bio3, bio4, bio12, bio18) and batch as fixed effects and the random effects listed above. *p*-values for climate terms were BH-corrected (batch excluded).

#### Life history associations

To compare life histories, we separated annual and perennial data and removed stolons (a confounding variable). We then ran the same LASSO analyses with 16 baseline traits as predictors of all four drought responses. Where results differed between life histories, we permuted life history labels 500 times to test whether differences in selection frequency exceeded those expected from sample size or power differences alone.

Apparent life history differences in climate associations were tested formally by adding a life history*climate interaction to the pooled models (see Climate–trait associations). Likewise, apparent differences in the stomatal conductance models were tested by adding life history interactions with the relevant terms to the pooled models, retaining all lower-order interactions and main effects. *P*-values for these interaction terms were BH-corrected within each trait across its tested climate variables.

## Results

### Drought Responses

Of the four drought traits, only wilt tolerance (i.e. time from wilting until senescence) was significantly associated with any climatic variable tested (Fig. 2A). Greater wilt tolerance was associated with higher isothermality (χ²(1) = 11.39, *p* < 0.01; Fig. 2B, Table S4) found in more southern and coastal accessions (Fig. 2C). Wilt tolerance trajectories varied across accessions (Fig. S4), and more tolerant individuals tended to reside in highly dynamic, stressful environments: high alpine, coastal, and desert regions (Fig. S5). Both pre-wilt (χ²(1) = 21.67, *p* < 0.001; Fig. 2D, Table S5) and post-wilt (χ²(1) = 13.72, *p* = 0.001; Fig. S6, Table S6) accessions from seasonally stable thermal environments had higher overall g_s_ than accessions where seasonal temperatures fluctuate.

**Figure 2.**
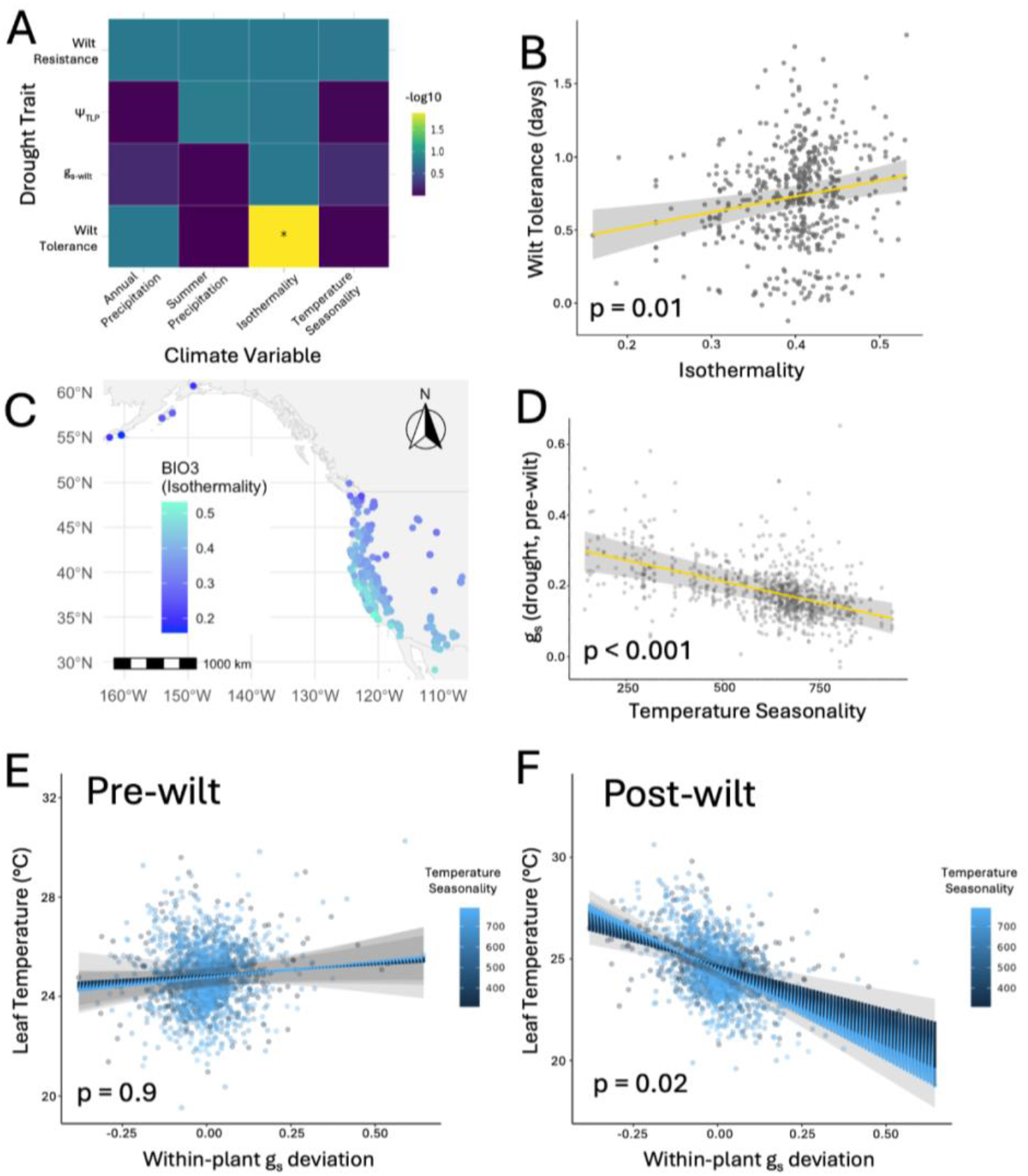
Climatic associations of drought response traits in *Mimulus guttatus.* A) ANOVA results of linear mixed effects models of four drought response traits (wilt time/resilience, turgor loss point, stomatal conductance at wilt, and death from wilt/wilt tolerance) as they relate to annual precipitation, summer precipitation, isothermality, and temperature seasonality. Colour and asterisks correspond to level of significance. B) Wilt tolerance in days and its relation with isothermality (*p* = 0.01). C) Map of accessions and where their climate falls along isothermality. D) Stomatal conductance during drought prior to wilting and its association with temperature seasonality (*p* < 0.001). The same relationship is observed post-wilt (Fig. S6). E) Relationship between stomatal conductance and leaf temperature during drought pre-wilt and its association with temperature seasonality (*p* = 0.9); compared to F) post-wilt (F; *p* = 0.02). Within-plant g_s_ deviation (from individual plant mean) is each time point difference in stomatal conductance relative to the individual plant’s mean stomatal conductance, which allows for comparison across plants. Leaf temperature measurements were collected over the course of two hours, during which the greenhouse likely experienced warming and increased solar radiation that affected increased T_leaf_ range.

We observed a phase shift in stomatal conductance before versus after wilt (g_s-drydown_*wilt_phase χ²(1) = 99.2, *p* < 0.001, Table S7): stomatal conductance and leaf temperature were not significantly associated before wilting (χ²(1) = 4.33, *p* > 0.05; Figs. 2E, S7 blue line) but were after wilting (g_s-postwilt_; Fig. S7 red line), which was also unchanged between life histories (Life History*g_s-drydown_*wilt_phase χ²(1) = 0.66, *p* = 0.46; Table S8, Fig. S7). This post-wilt association was also significantly affected by temperature seasonality (g_s-postwilt_*bio4 χ²(1) = 7.99, *p* = 0.02; Fig. 2F, Table S8). For the same change in conductance, accessions with high thermal seasonality had greater coupling of stomatal conductance and temperature (greater thermoregulation) than those from more thermally stable climates.

### Life History Drought Responses

We observed two differences between life histories when including life history as a covariate. We found that Ψ_TLP_ differed significantly between life histories: perennial Ψ_TLP_, in contrast to annuals, was associated with summer precipitation (Life history*bio18 χ²(1) = 7.86, *p* = 0.02; Fig. 3A, Table S9), where lower Ψ_TLP_ occurred where summers are dry (Fig. 3B). All plants exhibited greater rates of leaf cooling and g_s_ closer to wilt (Fig. S8), but only in annuals was it associated with drier summers (Life history*bio18*days to wilt χ²(1) = 9.09, *p* = 0.02, Fig. 3C, Table S10).

**Figure 3.**
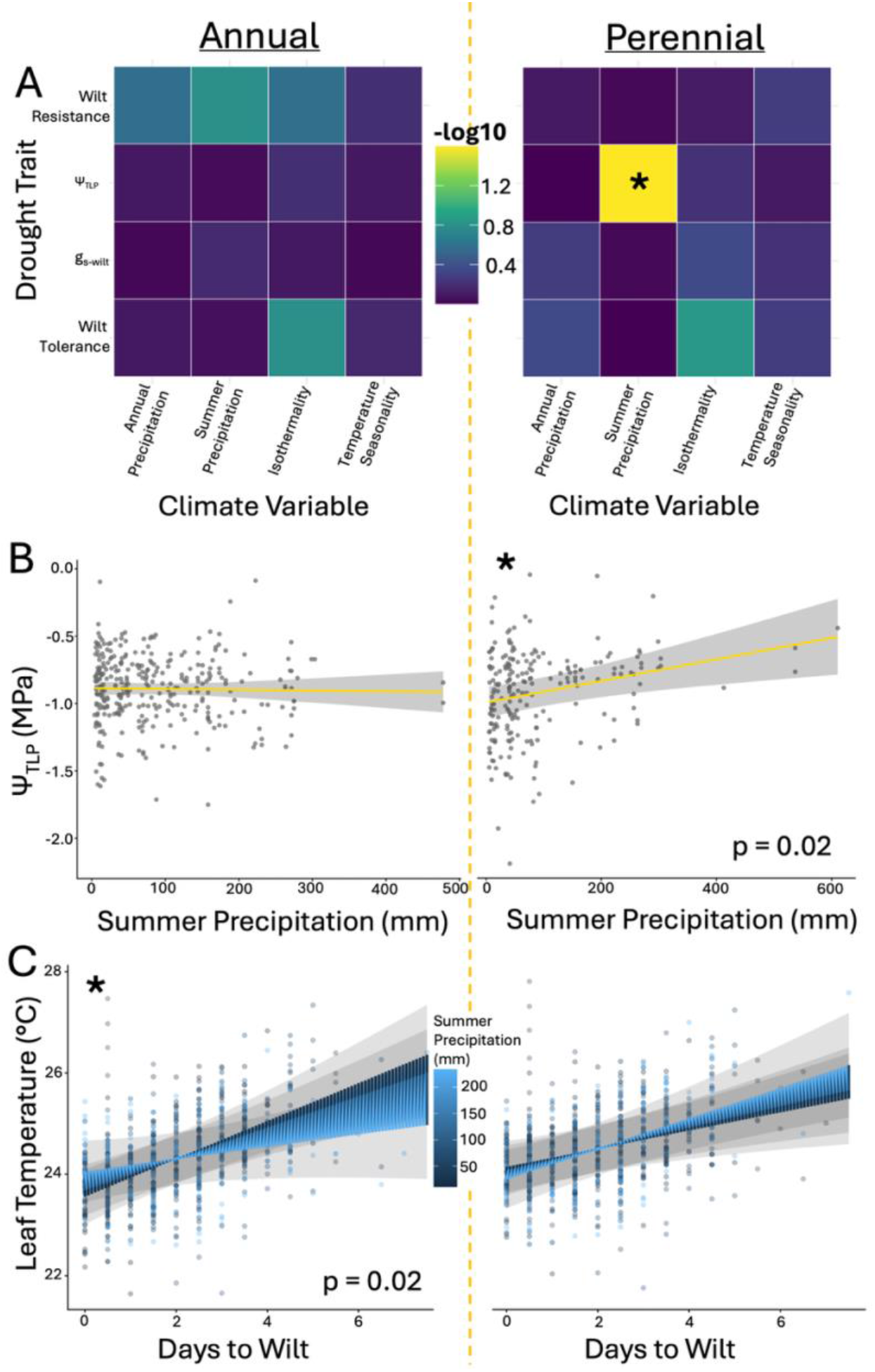
Differences in climatic associations and drought response across life histories (annual left, perennial right). A) ANOVA results of linear mixed effects models between annual and perennial accessions of four drought response traits (wilt time/resilience, turgor loss point, stomatal conductance at wilt, and death from wilt/wilt tolerance) as they relate to annual precipitation, summer precipitation, isothermality, and temperature seasonality. Colour and asterisks correspond to level of significance. B) The effect of summer precipitation on annual versus perennial turgor loss point (TLP; Life history*bio18 *p* = 0.02). C) The effect of summer precipitation on the relation between leaf temperature in the days prior to wilt in annual versus perennial accessions (Life history*bio18*days to wilt *p* = 0.02). Asterisks in B and C denote which life history shows significant association with parameter(s).

### Trait-trait Associations

None of the 17 baseline traits were selected as predictors of the four drought responses (Fig. S9). In contrast to g_s-wilt_, we found g_s-baseline_ complemented results from other studies and the leaf economic spectrum (Table S11). Higher g_s-baseline_ was associated with higher SPI (χ²(1) = 137.54, *p* < 0.001; Fig. 4B; Sack et al., 2003; Ocheltree et al., 2011) and lower δ^13^C (χ²(1) = 35.91, *p* < 0.01; Fig. 4C; Cernusak et al., 2013; Prieto et al., 2017). Additionally, we found higher g_s-baseline_ was associated with greater leaf lobing (χ²(1) = 25.10, *p* = 0.04; Fig. 4A) and later flowering date (χ²(1) = 71.57, p < 0.01; Fig. 4D). We also found that flowering date and δ^13^C in *M. guttatus* were negatively associated (χ²(1) = 20.65, p < 0.001; Fig. S10).

**Figure 4.**
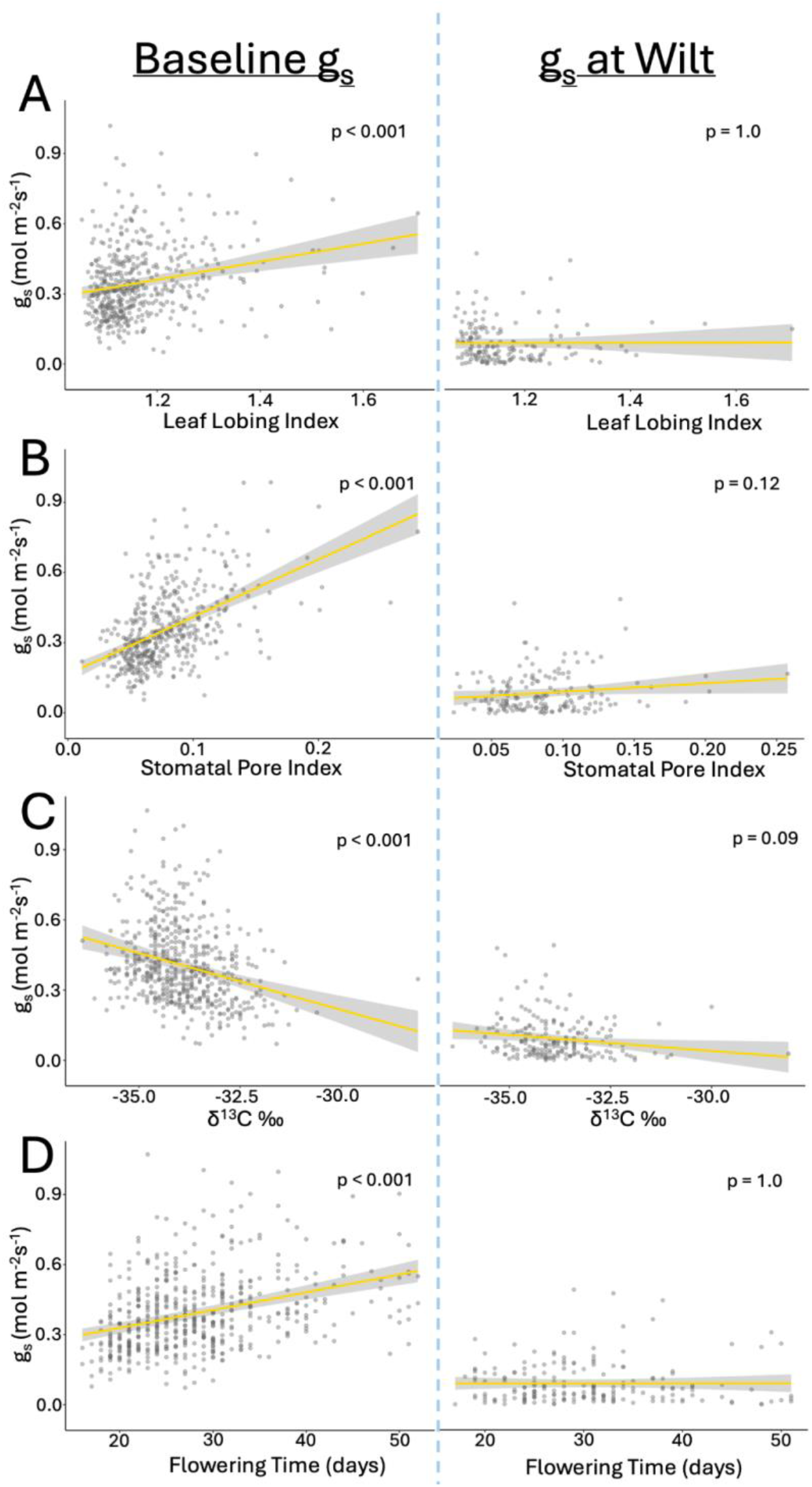
Stomatal conductance and its trait associations between baseline conductance and conductance at wilt. All associations were significant (*p* < 0.001) for baseline stomatal conductance but not for conductance at wilt. A) Leaf lobing; B) Stomatal pore index (ratio); which shows tremendous variation that is consistent when examining species across their range (Carlson et al., 2015); C) Carbon-13 isotope ratio; and D) Flowering time.

#### Climate-Trait Associations

Of the 17 baseline phenotypic traits, 13 were significantly associated with at least one of the four assessed climatic variables (*p* < 0.05; Fig. 5). Only four traits were associated with annual precipitation, all related to resource economics (specific leaf area χ²(610) = 10.89, *p* < 0.01; δ^15^N χ²(597) = 9.94, *p* < 0.01; δ^13^C χ²(593) = 14.23, *p* = 0.001; and stem thickness χ²(576) = 20.40, *p* < 0.001). Accessions from drier climates had thicker leaves and stems and higher δ^13^C (Mclean et al., 2013), associated with slower growing (Prieto et al., 2017), often drought-avoidant plants. Accessions also differed (regardless of life history) in nitrogen assimilation, with lower δ^15^N in plants that typically experienced drier summers but higher δ^15^N in plants that originate from overall drier climates. Summer precipitation also significantly affected SPI and trichome density: individuals from drier-summer habitats had lower SPI (χ²(474) = 28.22, *p* < 0.001) and more trichomes (χ²(484) = 20.63, *p* < 0.001).

**Figure 5.**
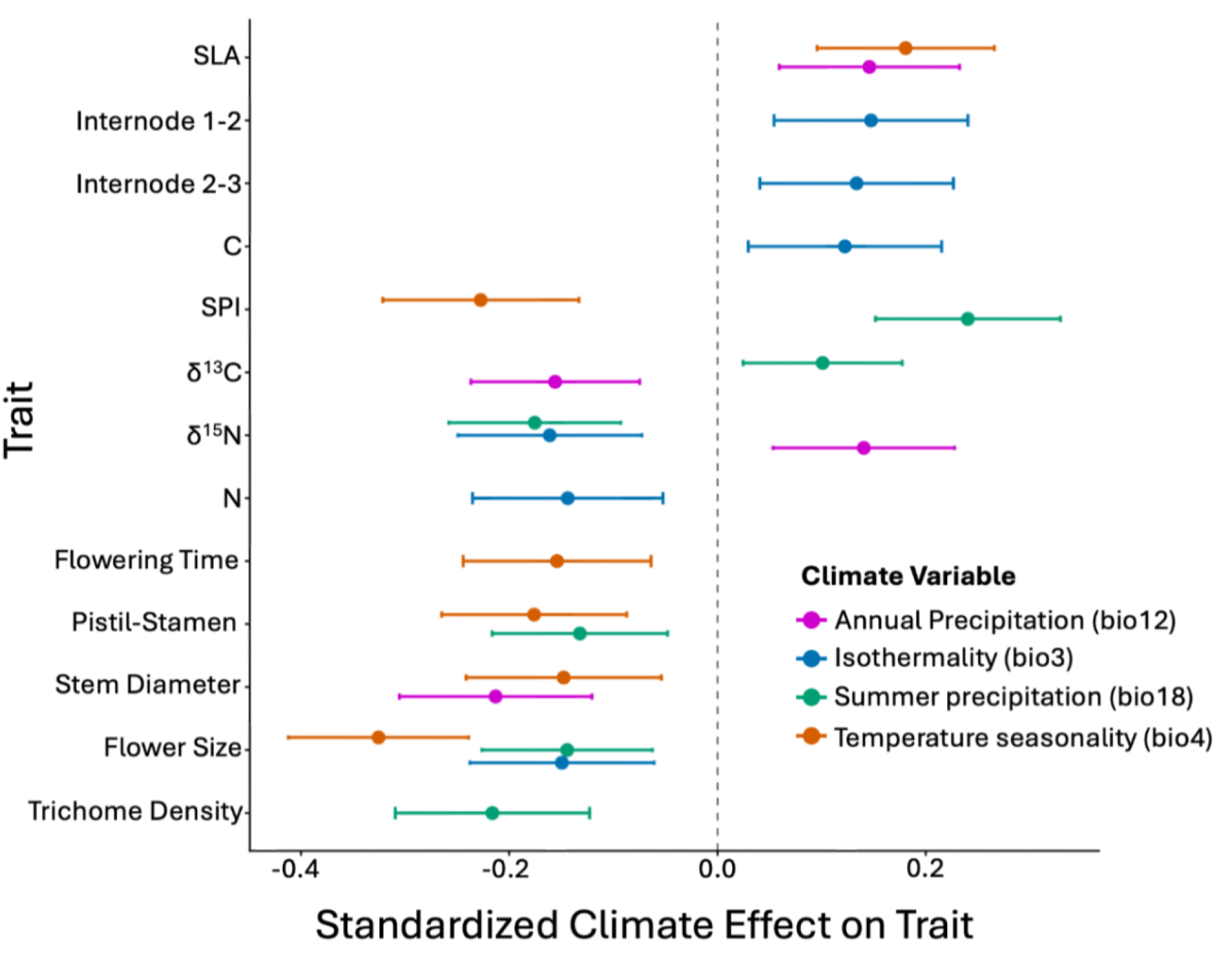
Phenotypic traits across the range of *Mimulus guttatus* and how they relate to climate. Location relative to 0.0 indicates the directionality of the association, either positive or negative. Colours indicate climatic variables: pink for annual precipitation, blue for isothermality, green for precipitation of the warmest quarter (summer precipitation), and orange for temperature seasonality. Confidence intervals are standard error bars.

Reproductive traits were more strongly associated with temperature (*p* < 0.05). Increased thermal variability (either diurnal or seasonal) and reduced summer precipitation were associated with reduced floral size (isothermality χ²(607) = 10.99, *p* < 0.01; seasonality χ²(505) = 54.18, *p* < 0.001; summer precip χ²(605) = 11.96, *p* < 0.01), and reduced pistil-stamen ratio (seasonality χ²(611) = 15.12, *p* = 0.001 ; summer precip χ²(613) = 9.48, *p* < 0.01), increased internode space (associated with bolting; isothermality χ²(603) = 9.63, *p* < 0.01 and χ²(600) = 7.93, *p* = 0.02 for internode 1-2 and 2-3 respectively), and earlier flowering time (seasonality χ²(588) = 11.22, p < 0.01).

## Discussion

We find a lack of immediate response to drought in *Mimulus guttatus* across its range and life histories. Contrary to expectations (Klein, 2014), we found no significant stomatal regulation following drought exposure prior to turgor loss. However, wilt tolerance (length of survival following wilt) varied with climate across the range. Accessions from climates with narrow daily but broad seasonal temperature ranges (e.g., high latitude) senesced faster following wilt, whereas those from climates with low seasonal relative to daily variation (e.g., coasts and deserts) survived longer (Fig. 2C). In perennials, accessions from dry-summer regions tolerated more negative water potentials before losing turgor (lower Ψ_TLP_; Fig. 3B), a relationship not significant in annuals. Finally, no baseline traits predicted drought response, suggesting phenology, morphology, and physiology under well-watered conditions are decoupled from physiological performance under stress. These results highlight the vulnerability of riparian plants to climatic change and the difficulty of predicting their responses, and show that a resource-acquisitive species can vary in its capacity for drought tolerance.

### Climatic association of wilt tolerance

We find isothermality is the only climate variable associated with any drought trait: wilt tolerance (Fig. 2B). As hypothesized, coastal and desert accessions survived more than two days longer after turgor loss than those from Alaska and the Pacific Northwest, consistent with local adaptation in this climatically broad species (Da Re et al., 2020). Under severe dehydration, embolisms (air bubbles in the xylem) restrict water transport (Kaack et al., 2021), and tension in the water column causes widespread cavitation and cellular collapse (Pereira et al., 2023). Species that risk embolism generally have mechanisms to prevent cavitation and to repair damage and recover (Bartlett et al., 2016). We did not measure water potential during the dry-down or secondary metabolites, so we lack direct measures of cavitation or its prevention. However, *M. guttatus* appears to acclimate to lower water potentials over repeated droughts (pers. obs.), implying a labile stress response possibly mediated by secondary metabolites that aid recovery and tolerance. Moreover, herbaceous species from Mediterranean climates with cyclical drought, which some *M. guttatus* populations experience, recover from dehydration better than species from more stable environments (Chaves et al., 2002; David et al., 1998).

We also found two key climate-associated differences between life histories. As predicted, perennials from regions with drier growing seasons tolerated more negative water potentials before losing turgor (Fig. 3B), possibly an evolutionary response for dormancy or slow growth through terminal drought or frost. However, we note that we cannot distinguish selection from other causes of clinal differences in perennials without extensive field experiments. In contrast, annuals from drier-summer climates showed stronger leaf cooling as they approached wilting (Fig. 3C), likely because their stomata remained fully open. Sustained high carbon assimilation is a common escape strategy for reproducing before terminal drought causes senescence. Thus, although neither life history showed stomatal regulation, perennials varied in drought tolerance and annuals in drought escape.

### Lack of stomatal response to drought

Before wilting, *M. guttatus* stomata did not behave as expected. Plants sensing water stress generally close their stomata before internal water potential changes substantially (Pirasteh-Anosheh et al., 2016; Scoffoni et al., 2017). The most cited mechanism is abscisic acid (ABA) production in roots and leaves, which triggers closure (McAdam et al., 2016; Buckley, 2019). Upon drying, *Arabidopsis thaliana* exhibited altered stomatal aperture within 10 minutes and near full closure within 45 minutes, a response attributed to ABA (Christmann et al., 2007). This rapid response maintains turgor and conserves water and is often associated with drought avoidance.

We found no such response, and our related work shows that *M. guttatus* stomata are unresponsive to ABA and other environmental stimuli (Tonet et al., in prep). While other studies have found differences in ABA responsiveness (Huber et al., 2019; Cardoso et al., 2020), our results show non-responsiveness across the range and both life histories. Our pre- and post-wilt leaf temperature–conductance relationships exemplify this lack of regulation. Stomatal regulation is closely tied to thermoregulation: open stomata cool leaves through transpiration (latent heat loss), whereas closed stomata raise leaf temperature relative to air (Buckley, 2019; Deva et al., 2020). Across all accessions, leaf temperature did not change before wilting (Fig. 2E) but increased steeply following wilt (Figs. 2F, S6). This suggests that post-wilt “stomatal conductance” reflects passive transpiration accompanying leaf-wide turgor loss and cellular collapse, not stomatal regulation (Brodribb, 2003). In other species, the pre- to post-wilt phase shift often marks a switch from risky to conservative water use. Across ∼100 woody species, behaviour before turgor loss varied widely, but beyond Ψ_TLP_ nearly all conserved water to limit embolism (Martin-StPaul et al., 2017). Although stomatal conductance at wilting averaged 74% below baseline, this decline took days, and its association with wilting strengthened as plants approached wilt (Fig. S9). Thus, we found no evidence that *M. guttatus* ever shifted to water conservation.

Importantly, our results indicate a lack of stomatal response, not stomatal function. We observed fluctuations in g_s-drydown_ between morning and afternoon and across days (Fig. S4), although the mechanism regulating aperture is unclear. Although species vary in stomatal responsiveness to drought, from water conservation to risk-taking (Bartlett et al., 2016; Martin-StPaul et al., 2017), to our knowledge this is the first study to document such an extreme lack of stomatal response.

### Trait-trait associations are flexible

Given these results, it is unsurprising that g_s-wilt_ was not associated with any baseline trait. In contrast, gs-baseline was significantly associated with leaf lobing, stomatal pore index, δ^13^C, and flowering date. This loss of associations from gs-baseline to gs-wilt further indicates a shift under drought, especially as two of them (SPI and δ^13^C) followed expected relationships with gs-baseline. Greater SPI increases conductance, and higher conductance typically increases carbon gain and discrimination against ^13^C, lowering δ^13^C. Leaf lobing, however, was positively related to gs-baseline, contrary to our expectation. Because lobing increases turbulence and convective cooling, reducing the need for transpiration, we predicted plants from hot, dry regions would have more lobed leaves and lower water-use (Vogel, 2009). This reversal *in M. guttatus* may reflect trichome presence, which increases where summers are dry both in the current study as well as other *Mimulus* species (Fig. 5; see Branch et al., 2024b), corroborating expectations for mesic versus xeric climates (Galdon-Armero et al., 2018; Branch et al., 2024b). These trichomes could alter the boundary layer, potentially making lobing less effective or less necessary.

Flowering time appears more labile than the other traits. Higher conductance is usually associated with rapid growth and early flowering (Chaves et a., 2003; Kooyers, 2015), yet here we find higher gs-baseline associated with later flowering. Similarly, flowering time and δ^13^C are usually positively correlated, with early flowering and low δ^13^C indicating escape and late flowering and high δ^13^C indicating avoidance, as in *Arabidopsis* (Kooyers et al., 2015) and *Conyza canadensis* (Nagy et al., 2024). In contrast, later-flowering plants had lower leaf δ^13^C (Fig. S10), matching annual *M. guttatus* (Kooyers et al., 2015), in which larger plants generally flowered later without affecting δ^13^C. We did not measure total biomass, but our size proxies (stem thickness and preliminary leaf mass and size data) were unrelated to stomatal conductance, although such relationships are plausible (Díaz et al., 2016).

### Climatic association of traits does not predict drought response

Across its range, *M. guttatus* generally presents escape-type traits, to an extent that varies with climate. Accessions from more thermally variable environments tend to have cheaper, broader, thinner leaves, rapid growth (bolting), and earlier reproduction with smaller, likely more selfing flowers (Fig. 5), consistent with drought escape. More resource-acquisitive (escape) accessions also occurred where annual precipitation was higher, with broader leaves, higher δ^15^N, thinner stems, and lower δ^13^C. In contrast, accessions from less thermally variable and drier climates had avoidance-type traits: thicker leaves and stems, higher δ^13^C, and later flowering. While higher δ^13^C indicates greater intrinsic water-use efficiency (Spasojevic & Weber, 2021) and accessions from drier climates had higher δ^13^C (Fig. 5), the range of δ^13^C across accession (−35‰ to −31‰) is narrow relative to natural variation across the Plant Kingdom and may have little effect on water-use efficiency. Because δ^13^C often reflects stomatal regulation (Spasojevic & Weber, 2021), it is puzzling that this relationship persists when *M. guttatus* stomata do not close in response to drought.

Traits were also decoupled along precipitation gradients. δ^15^N depended on the precipitation variable, being lower in accessions from regions with more summer rainfall and less annual precipitation. These differences may reflect range-wide heritable variation in nitrogen processing, possibly including local adaptation to nitrogen availability. Indeed, adaptations that equalize nitrogen acquisition across diverse environments may appear as isotopic differences in a common greenhouse (Spasojevic & Weber, 2021).

Overall, distinct trait combinations in *M. guttatus* were associated with different climates. Resource economic traits tracked annual precipitation, which suggests plant acquisitiveness is strongly associated with precipitation (Prieto et al., 2017). Water availability shapes carbon and nitrogen uptake, thus range-wide precipitation patterns could result in differing acquisition physiologies that persist in a common greenhouse (Spasojevic & Weber, 2021). Reproductive traits, in contrast, tracked temperature more than precipitation, differing in flower size, phenology, and potential for selfing. These could reflect the thermal sensitivity of reproduction (Sage et al., 2012; Branch & Sage, 2018; Arathi & Smith, 2023).

## Conclusions

While the mechanisms underlying variation in wilt tolerance and the lack of stomatal regulation are unknown, our experiment yields two conclusions concerning vulnerability to global change and nuance to the leaf economic spectrum. *Mimulus guttatus* is an herbaceous riparian plant generally characterized as a drought escaper. Under simulated terminal drought, we observed a lack of water conservation. Although escape species generally do not respond immediately to drought, the absence of stomatal regulation as drought intensified is surprising. We did, however, observe that wilt tolerance varied with climate across accessions, suggesting past capacity to evolve traits of other drought strategies; future work should examine drought recovery and cavitation resistance. Tolerating dehydration after wilting helps only in acute, not prolonged, drought. Thus, *M. guttatus* appears to lack an effective drought response and is merely at the mercy of water availability and whether it has already reproduced. As hydrological change alters water access and shortens the growing season, drought-escaping species may become more vulnerable if drought occurs too early in the season. Our results highlight the vulnerability of ephemeral river species to global change and provide an exceptional example of range-wide absence of stomatal regulation.

## Data accessibility statement

Upon publication of this manuscript, all data will be available on *Dryad Digital Repository* and all coding available on *GitHub*.

## Statement of authorship

HB, PR, VT, HAC, CB, EE, and JC designed and implemented the experiment. HB conducted the bulk of the analyses, result interpretation, and wrote the manuscript. PR significantly contributed to the analysis and analytical design and result interpretation. HB, PR, and VT conducted all stomatal conductance and temperature measurements. HB was responsible for all other leaf characteristics (SLA, lobing, anatomical measurements, isotopic measurements), apart from TLP, which was measured by VT with help from HB and PR. HAC was responsible for the refresher generation, with help from HB, PR, VT, and JC. HAC was responsible for floral characteristics with help from PR. AF, LH, AS, EE, and JC collected *in situ* data and samples for later processing. DL and HB processed leaf lobing index. MV-M, KT, BB, JC conducted field collections of seeds. HB was responsible for all data management. All authors contributed to manuscript editing.

## Acknowledgements

We are grateful to Christopher Bolick and Nathan Guzzo for helping with plant care in the Yale Plant Growth Facility and Brad Erkkila at the Yale Analytical and Stable Isotope Center for assistance with stable isotope mass spectrometry. We thank Pei-Wei Sun for assistance in species niche modeling, Keric Lamb for providing the code to process leaf margins and lobing percentage, and Lila Fishman, Paige Smith, and David Lowry for providing accessions. We thank Ken Thompson for assistance with initial fieldwork collection. We thank France DuVal for assistance in weighing leaf material for isotope measurements. HB was supported by a NSF PRFB 2305993 and NSERC PDF 578168 and PR, VT, and HAC were supported by the Yale Institute for Biospheric Studies through G. Evelyn Hutchinson Environmental Postdoctoral Fellowships. AF and AS were supported by the Yale Institute for Biospheric Studies Summer Undergraduate Research Experience program, DL was supported by Yale’s Summer Undergraduate Research Fellowship, and LH was supported by a Yale College First-Year Summer Research Fellowship.

### Box 1. Definitions of Terms

Terminal drought – Water availability that decreases continuously; drying of an ephemeral river

Baseline – Phenotypes characterized during benign conditions

Wilt Resistance – Time to wilt from dry-down treatment initiation

Wilt Tolerance – Length of survival following wilt

g_s-baseline_ – Stomatal conductance prior to drought

g_s-wilt_ – Stomatal conductance at the time wilt occurs

g_s-drydown_ – Timeseries of stomatal conductance throughout the dry-down treatment

g_s-prewilt_ – Timeseries of stomatal conductance during dry-down but prior to wilting

g_s-postwilt_ – Timeseries of stomatal conductance during dry-down following wilt

