## Supplemental Material for "Divergence in wilt tolerance across the range of *Mimulus guttatus* in the absence of stomatal regulation"

**Tables**

**Table S1.** Maximum entropy species distribution modeling of 3189 GBIF collected locations of *Mimulus guttatus.*

| **Variable** | **Percent Contribution** | **Permutation Importance** |
| --- | --- | --- |
| **bio4** | **27.5** | **14.8** |
| **bio3** | **26** | **48.2** |
| **bio18** | **19.3** | **2.2** |
| bio8 | 10.2 | 3.3 |
| bio11 | 4 | 2.2 |
| bio19 | 3.9 | 15 |
| bio6 | 3.6 | 0 |
| bio7 | 1.8 | 0 |
| bio14 | 0.9 | 6.9 |
| bio9 | 0.8 | 0.8 |
| bio10 | 0.6 | 0.9 |
| bio5 | 0.4 | 1.3 |
| bio12 | 0.3 | 0.5 |
| bio2 | 0.2 | 1 |
| bio15 | 0.2 | 0.8 |
| bio1 | 0.2 | 1.6 |
| bio13 | 0.1 | 0 |
| bio16 | 0 | 0 |
| bio17 | 0 | 0.4 |

**Table S2.** Level of missing data for each trait in order from most to least in a dataset of 658 samples of *Mimulus guttatus*.

| **Trait** | **n** | **Missing** | **Percent missing** |
| --- | --- | --- | --- |
| *g_s-wilt_ | 222 | 436 | 66.3 |
| Wilt resistance | 471 | 187 | 28.4 |
| SPI | 484 | 174 | 26.4 |
| Trichome Density | 491 | 167 | 25.4 |
| Lobing | 495 | 163 | 24.8 |
| Wilt Tolerance | 520 | 138 | 21 |
| Turgor Loss Point | 557 | 101 | 15.3 |
| UV Proportion | 567 | 91 | 13.8 |
| Stem Diameter | 584 | 74 | 11.2 |
| gs | 601 | 57 | 8.7 |
| C | 602 | 56 | 8.5 |
| CN | 602 | 56 | 8.5 |
| 𝛿^15^N | 603 | 55 | 8.4 |
| 𝛿^13^C | 603 | 55 | 8.4 |
| N | 603 | 55 | 8.4 |
| Internode12 | 610 | 48 | 7.3 |
| Internode23 | 610 | 48 | 7.3 |
| Node | 615 | 43 | 6.5 |
| Pistil-Stamen | 620 | 38 | 5.8 |
| Specific Leaf Area | 621 | 37 | 5.6 |
| Pistil Length | 623 | 35 | 5.3 |
| Corolla Tube Length | 624 | 34 | 5.2 |
| Corolla Width | 625 | 33 | 5 |
| Stamen Length | 625 | 33 | 5 |
| Total Flower Length | 626 | 32 | 4.9 |
| Flowering Date | 631 | 27 | 4.1 |
| Stolons | 631 | 27 | 4.1 |

^*^Stomatal conductance of wilt was measured on only a subset of the data

**Table S3.** Transformations used for each LASSO model.

| **Trait** | **N** | **Transform** | **Shapiro-Wilk W** | **Shapiro-Wilk p** | **Skewness** |
| --- | --- | --- | --- | --- | --- |
| Wilt Resistance | 206 | none | 0.99 | 0.162 | 0.311 |
| Wilt Tolerance | 222 | log(x+1) | 0.991 | 0.158 | -0.145 |
| g_s-wilt_ | 91 | sqrt | 0.979 | 0.156 | 0.356 |
| TLP | 230 | none | 0.965 | 0 | -0.742 |
| g_s-baseline_ | 262 | sqrt | 0.99 | 0.058 | 0.214 |

**Table S4.** ANOVA Type III results for climate-drought response associations.

| **Trait** | **Climate** | **Chisq** | **Df** | **p_value** | **p_adj** |
| --- | --- | --- | --- | --- | --- |
| Wilt resistance | bio3_z | 3.185 | 1 | 0.074 | 0.17 |
| Wilt resistance | bio4_z | 2.738 | 1 | 0.098 | 0.174 |
| Wilt resistance | bio18_z | 3.453 | 1 | 0.063 | 0.17 |
| Wilt resistance | bio12_z | 3.241 | 1 | 0.072 | 0.17 |
| Wilt tolerance | bio3_z | 11.39 | 1 | 0.001 | 0.012 |
| Wilt tolerance | bio4_z | 0.032 | 1 | 0.858 | 0.982 |
| Wilt tolerance | bio18_z | 0 | 1 | 0.997 | 0.997 |
| Wilt tolerance | bio12_z | 3.608 | 1 | 0.057 | 0.17 |
| gs-wilt | bio3_z | 3.333 | 1 | 0.068 | 0.17 |
| gs-wilt | bio4_z | 0.805 | 1 | 0.37 | 0.591 |
| gs-wilt | bio18_z | 0.01 | 1 | 0.92 | 0.982 |
| gs-wilt | bio12_z | 0.637 | 1 | 0.425 | 0.618 |
| TLP | bio3_z | 2.843 | 1 | 0.092 | 0.174 |
| TLP | bio4_z | 0.135 | 1 | 0.714 | 0.952 |
| TLP | bio18_z | 5.444 | 1 | 0.02 | 0.157 |
| TLP | bio12_z | 0.022 | 1 | 0.882 | 0.982 |

**Table S5.** ANOVA Type III results of climatic associations with stomatal conductance during drought but prior to wilt.

| **term** | **Chisq** | **Df** | **Pr(>Chisq)** | **p_adj** |
| --- | --- | --- | --- | --- |
| (Intercept) | 122.046 | 1 | 0 | 0 |
| Batch | 14.844 | 1 | 0 | 0 |
| bio3_z | 0.011 | 1 | 0.915 | 0.915 |
| bio4_z | 21.669 | 1 | 0 | 0 |
| bio12_z | 0.166 | 1 | 0.683 | 0.911 |
| bio18_z | 1.324 | 1 | 0.25 | 0.5 |
| time_to_wilt | 5.042 | 1 | 0.025 | 0.059 |
| tp | 24.322 | 1 | 0 | 0 |
| time_to_wilt:bio3_z | 0.093 | 1 | 0.76 | 0.912 |
| time_to_wilt:bio4_z | 0.201 | 1 | 0.654 | 0.911 |
| time_to_wilt:bio12_z | 0.934 | 1 | 0.334 | 0.572 |
| time_to_wilt:bio18_z | 0.038 | 1 | 0.845 | 0.915 |

**Table S6.** ANOVA Type III results of climatic associations with stomatal conductance after wilt.

| **term** | **Chisq** | **Df** | **Pr(>Chisq)** | **p_adj** |
| --- | --- | --- | --- | --- |
| (Intercept) | 62.952 | 1 | 0 | 0 |
| Batch | 2.809 | 1 | 0.094 | 0.187 |
| bio3_z | 4.66 | 1 | 0.031 | 0.093 |
| bio4_z | 13.72 | 1 | 0 | 0.001 |
| bio12_z | 0.021 | 1 | 0.885 | 0.885 |
| bio18_z | 0.731 | 1 | 0.392 | 0.562 |
| time_since_wilt | 12.466 | 1 | 0 | 0.002 |
| tp | 0.647 | 1 | 0.421 | 0.562 |
| time_since_wilt:bio3_z | 1.631 | 1 | 0.202 | 0.346 |
| time_since_wilt:bio4_z | 3.736 | 1 | 0.053 | 0.128 |
| time_since_wilt:bio12_z | 0.245 | 1 | 0.621 | 0.737 |
| time_since_wilt:bio18_z | 0.176 | 1 | 0.675 | 0.737 |

**Table S7.** ANOVA Type III results of the association of stomatal conductance prior to and following wilt and its impact on leaf temperature.

| **term** | **Chisq** | **Df** | **Pr(>Chisq)** | **p_adj** |
| --- | --- | --- | --- | --- |
| (Intercept) | 7224.455 | 1 | 0 | 0 |
| Batch | 143.832 | 1 | 0 | 0 |
| gs_within | 2.212 | 1 | 0.137 | 0.137 |
| post_wilt | 7.159 | 1 | 0.007 | 0.009 |
| gs_mean | 35.011 | 1 | 0 | 0 |
| tp | 204.859 | 1 | 0 | 0 |
| gs_within:post_wilt | 99.209 | 1 | 0 | 0 |

**Table S8.** ANOVA Type III results of the life history differences in association of stomatal conductance prior to and following wilt and its impact on leaf temperature.

| **term** | **Chisq** | **Df** | **Pr(>Chisq)** | **p_adj** |
| --- | --- | --- | --- | --- |
| (Intercept) | 6775.004 | 1 | 0 | 0 |
| Batch | 149.423 | 1 | 0 | 0 |
| LifeHistory | 3.648 | 1 | 0.056 | 0.088 |
| gs_within | 2.662 | 1 | 0.103 | 0.141 |
| post_wilt | 5.848 | 1 | 0.016 | 0.029 |
| gs_mean | 36.252 | 1 | 0 | 0 |
| tp | 208.49 | 1 | 0 | 0 |
| LifeHistory:gs_within | 0.694 | 1 | 0.405 | 0.46 |
| LifeHistory:post_wilt | 0.022 | 1 | 0.882 | 0.882 |
| gs_within:post_wilt | 51.191 | 1 | 0 | 0 |
| LifeHistory:gs_within:post_wilt | 0.655 | 1 | 0.418 | 0.46 |

**Table S9.** ANOVA Type III results for the climate associations in turgor loss point that differ between life histories.

| **term** | **Chisq** | **Df** | **p_value** | **p_adj** |
| --- | --- | --- | --- | --- |
| LifeHistory:bio18_z | 7.859 | 1 | 0.005 | 0.02 |
| LifeHistory:bio3_z | 0.084 | 1 | 0.772 | 0.772 |
| LifeHistory:bio4_z | 0.472 | 1 | 0.492 | 0.772 |
| LifeHistory:bio12_z | 0.117 | 1 | 0.733 | 0.772 |

**Table S10.** ANOVA Type III results for life history differences in how climate interacts with stomatal conductance following wilt to affect leaf cooling.

| **term** | **Chisq** | **Df** | **Pr(>Chisq)** | **p_adj** |
| --- | --- | --- | --- | --- |
| (Intercept) | 2368.76 | 1 | 0 | 0 |
| Batch | 197.671 | 1 | 0 | 0 |
| LifeHistory | 4.901 | 1 | 0.027 | 0.125 |
| gs_within | 0.822 | 1 | 0.364 | 0.491 |
| time_to_wilt | 16.271 | 1 | 0 | 0.001 |
| bio3_z | 3.466 | 1 | 0.063 | 0.157 |
| bio4_z | 3.854 | 1 | 0.05 | 0.145 |
| bio12_z | 1.118 | 1 | 0.29 | 0.423 |
| bio18_z | 3.346 | 1 | 0.067 | 0.157 |
| gs_mean | 4.895 | 1 | 0.027 | 0.125 |
| tp | 0.003 | 1 | 0.956 | 0.96 |
| LifeHistory:gs_within | 1.592 | 1 | 0.207 | 0.345 |
| LifeHistory:time_to_wilt | 2.617 | 1 | 0.106 | 0.231 |
| gs_within:time_to_wilt | 0.005 | 1 | 0.944 | 0.96 |
| gs_within:bio3_z | 0.153 | 1 | 0.695 | 0.842 |
| time_to_wilt:bio3_z | 0.973 | 1 | 0.324 | 0.453 |
| LifeHistory:bio3_z | 3.512 | 1 | 0.061 | 0.157 |
| gs_within:bio4_z | 0.003 | 1 | 0.96 | 0.96 |
| time_to_wilt:bio4_z | 4.346 | 1 | 0.037 | 0.13 |
| LifeHistory:bio4_z | 1.364 | 1 | 0.243 | 0.386 |
| gs_within:bio12_z | 0.005 | 1 | 0.942 | 0.96 |
| time_to_wilt:bio12_z | 2.168 | 1 | 0.141 | 0.274 |
| LifeHistory:bio12_z | 2.396 | 1 | 0.122 | 0.25 |
| gs_within:bio18_z | 0.018 | 1 | 0.893 | 0.96 |
| time_to_wilt:bio18_z | 6.495 | 1 | 0.011 | 0.076 |
| LifeHistory:bio18_z | 4.791 | 1 | 0.029 | 0.125 |
| LifeHistory:gs_within:time_to_wilt | 4.383 | 1 | 0.036 | 0.13 |
| LifeHistory:gs_within:bio3_z | 0.048 | 1 | 0.826 | 0.96 |
| LifeHistory:time_to_wilt:bio3_z | 1.28 | 1 | 0.258 | 0.392 |
| LifeHistory:gs_within:bio4_z | 0.151 | 1 | 0.698 | 0.842 |
| LifeHistory:time_to_wilt:bio4_z | 1.73 | 1 | 0.188 | 0.33 |
| LifeHistory:gs_within:bio12_z | 1.843 | 1 | 0.175 | 0.322 |
| LifeHistory:time_to_wilt:bio12_z | 4.01 | 1 | 0.045 | 0.144 |
| LifeHistory:gs_within:bio18_z | 0.474 | 1 | 0.491 | 0.637 |
| LifeHistory:time_to_wilt:bio18_z | 9.091 | 1 | 0.003 | 0.022 |

**Table S11.** ANOVA Type III results for baseline stomatal conductance and its association with four other baseline traits.

| **Term** | **Chisq** | **Df** | **Pr(>Chisq)** |
| --- | --- | --- | --- |
| Flowering | 71.566 | 1 | 0 |
| Lobing | 25.096 | 1 | 0 |
| SPI | 137.54 | 1 | 0 |
| d13C | 35.906 | 1 | 0 |

**Figures**

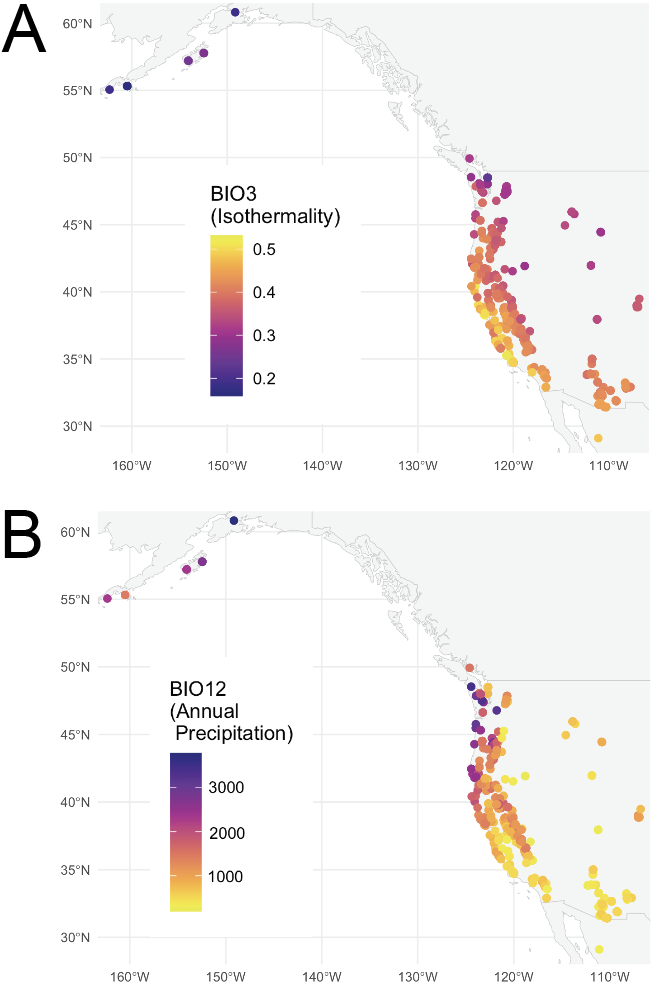

**Figure S1.** Annual precipitation levels across accessions.

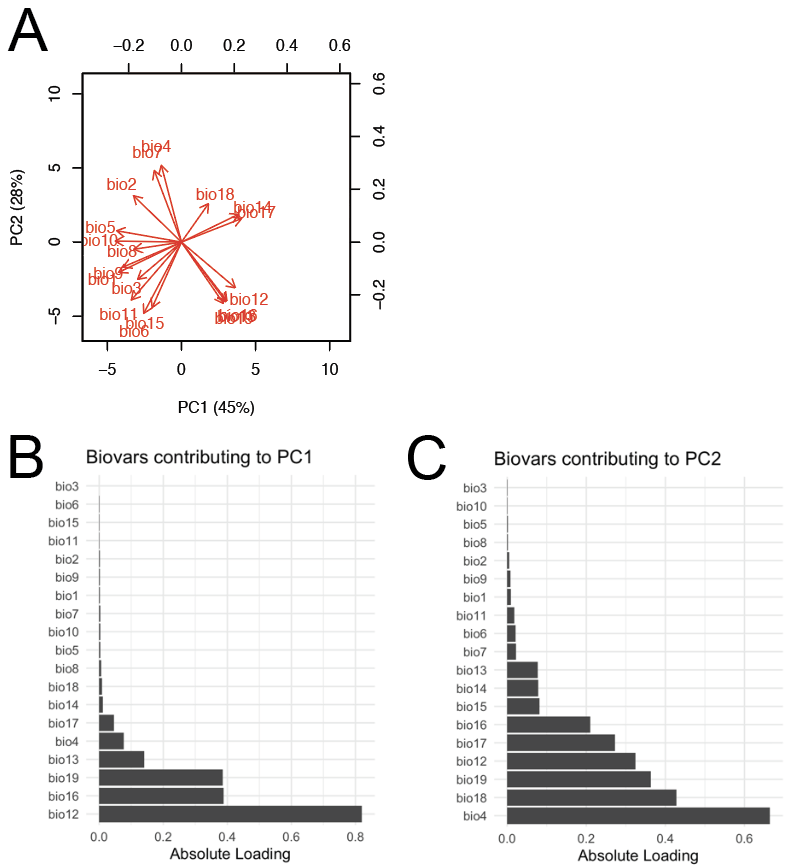

**Figure S2.** PCA of all biovars and their contributions to variation across 715 accessions of *Mimulus guttatus.* A) PCA. B) PC1 biovar contributions. C) PC2 biovar contributions.

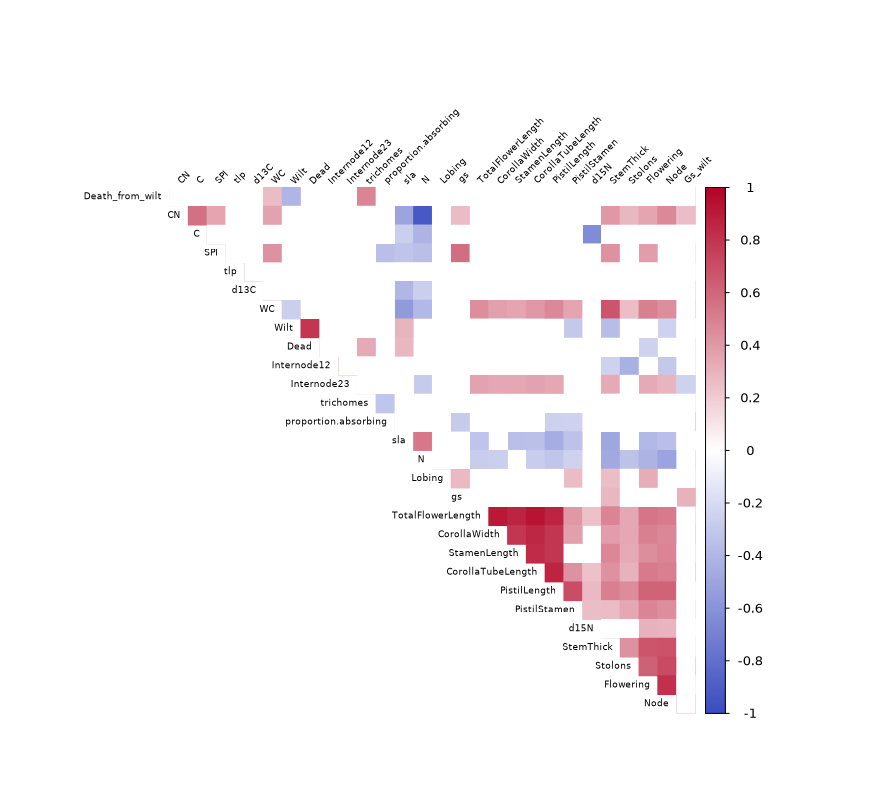

**Figure S3.** Correlation matrix of 28 traits.

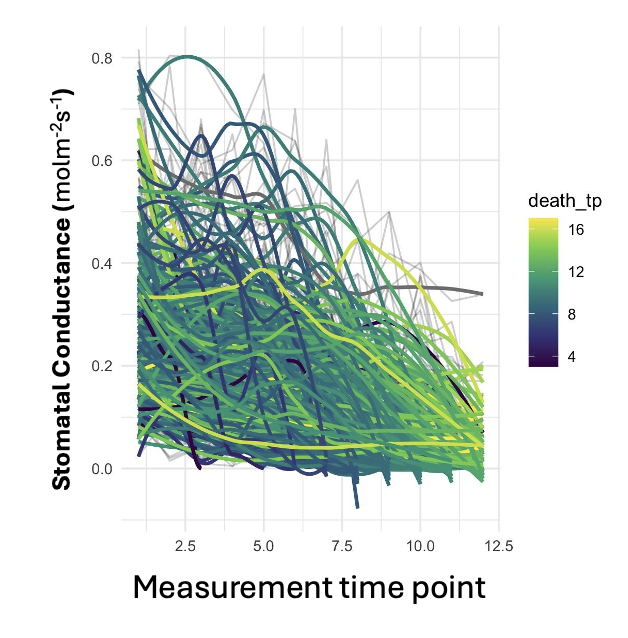

**Figure S4.** Stomatal conductance during dry-down of 222 *Mimulus guttatus* accessions. Measurements were taken twice a day and colours indicate the timing of death of each individual. Death_tp (death timepoint) thus indicates how many measurements were conducted prior to death (i.e., measurement 4 corresponds to 2 days of dry down treatment, while 16 indicates 8 days).

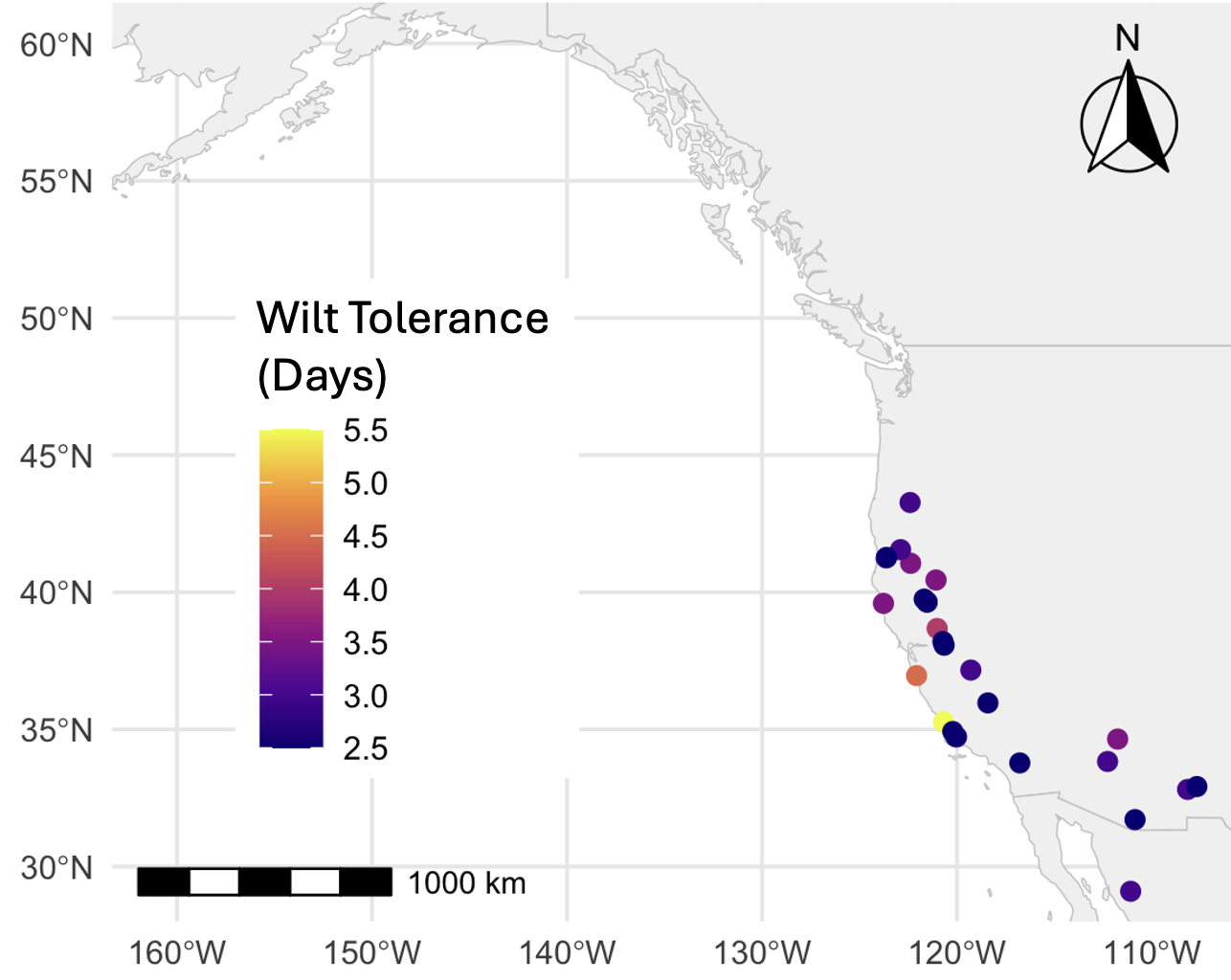

**Figure S5.** Map of wilt tolerance in days of accessions that withstood wilt for more than 2 days.

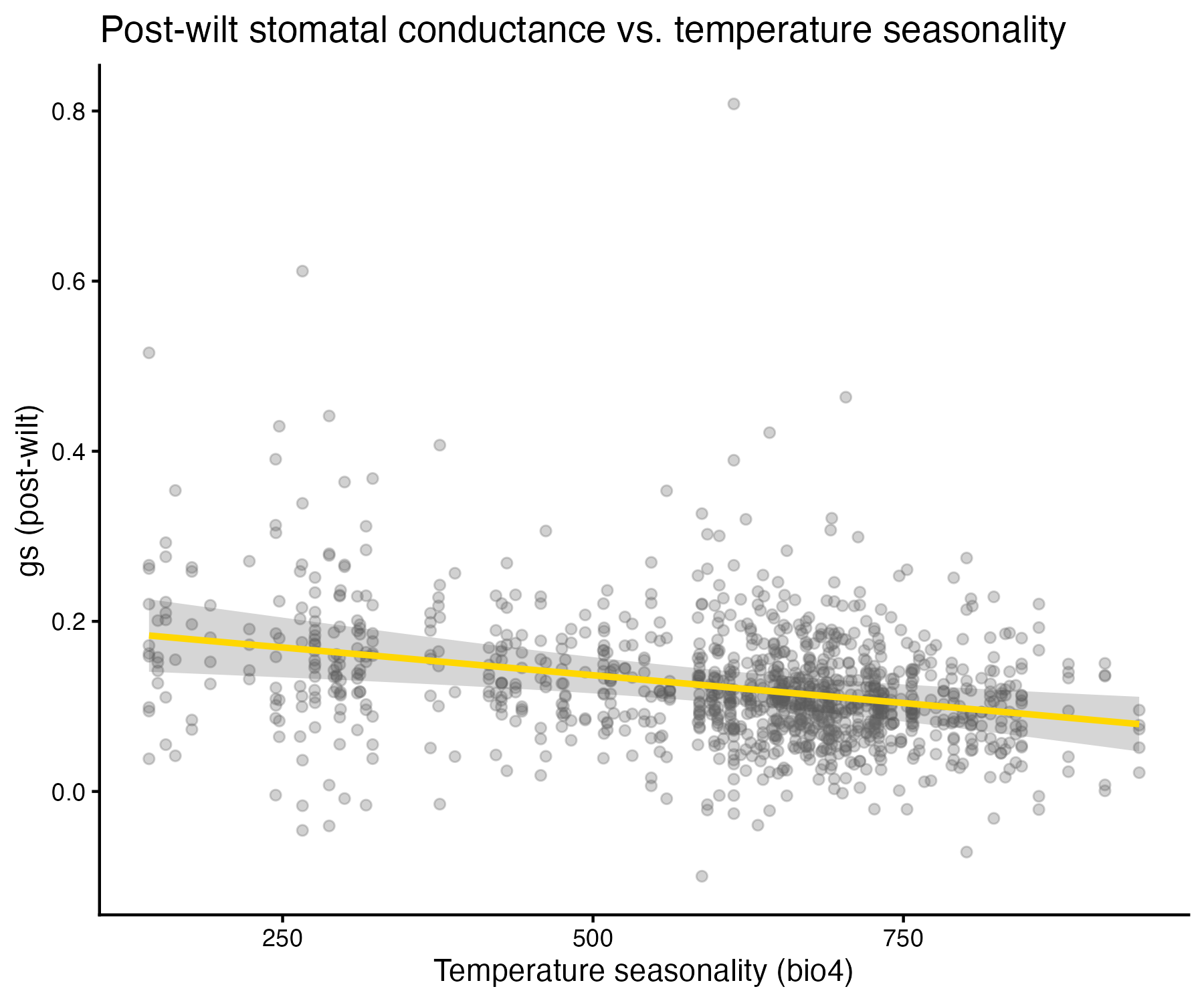

**Figure S6.** Relationship between stomatal conductance following wilt and temperature seasonality (*p* = 0.001).

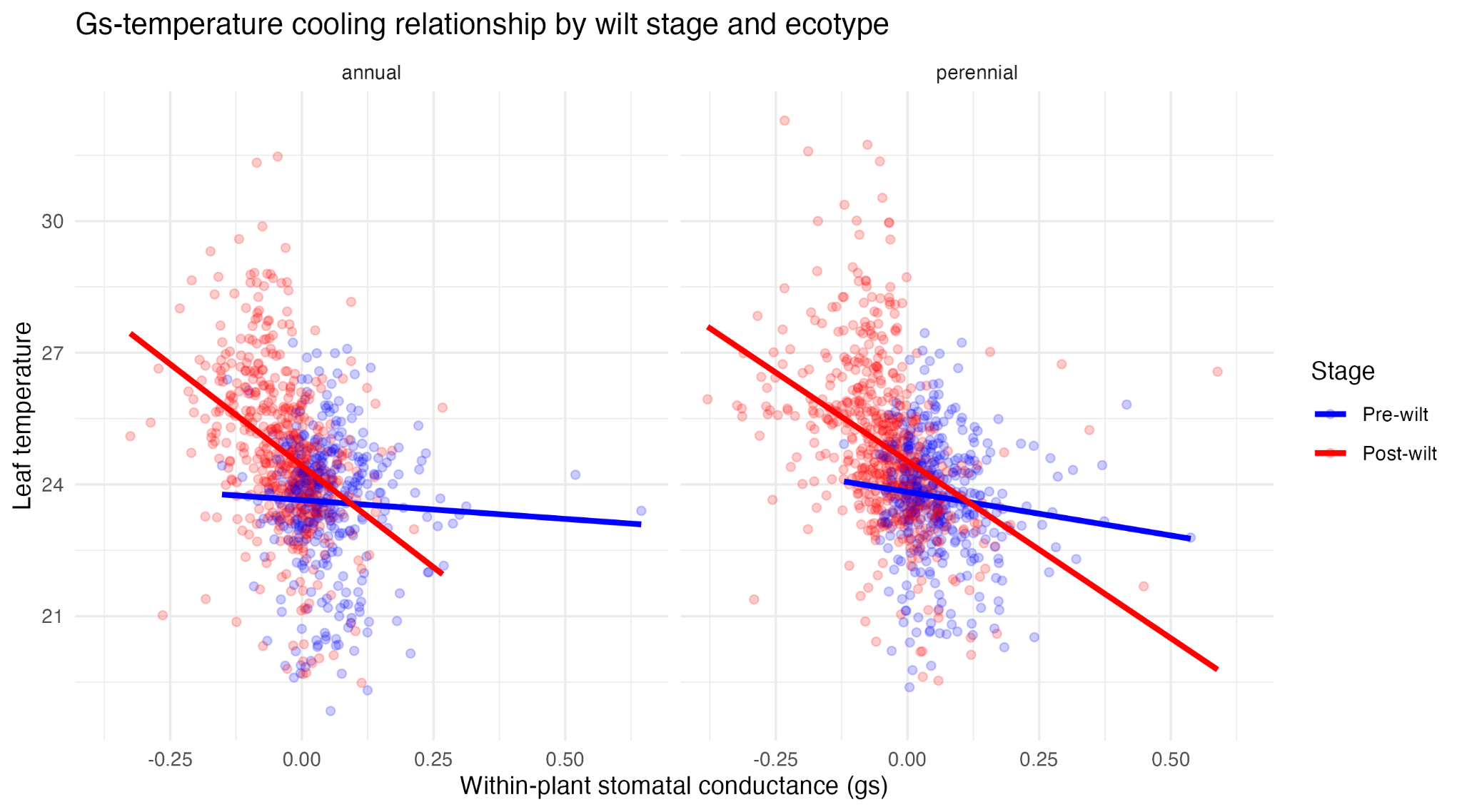

**Figure S7.** Relationship between leaf temperature and stomatal conductance during dry-down. Relationship is the same across annual and perennial (*p* = 0.46), and exhibit a phase-shift from pre-wilt to post-wilt (*p* < 0.001).

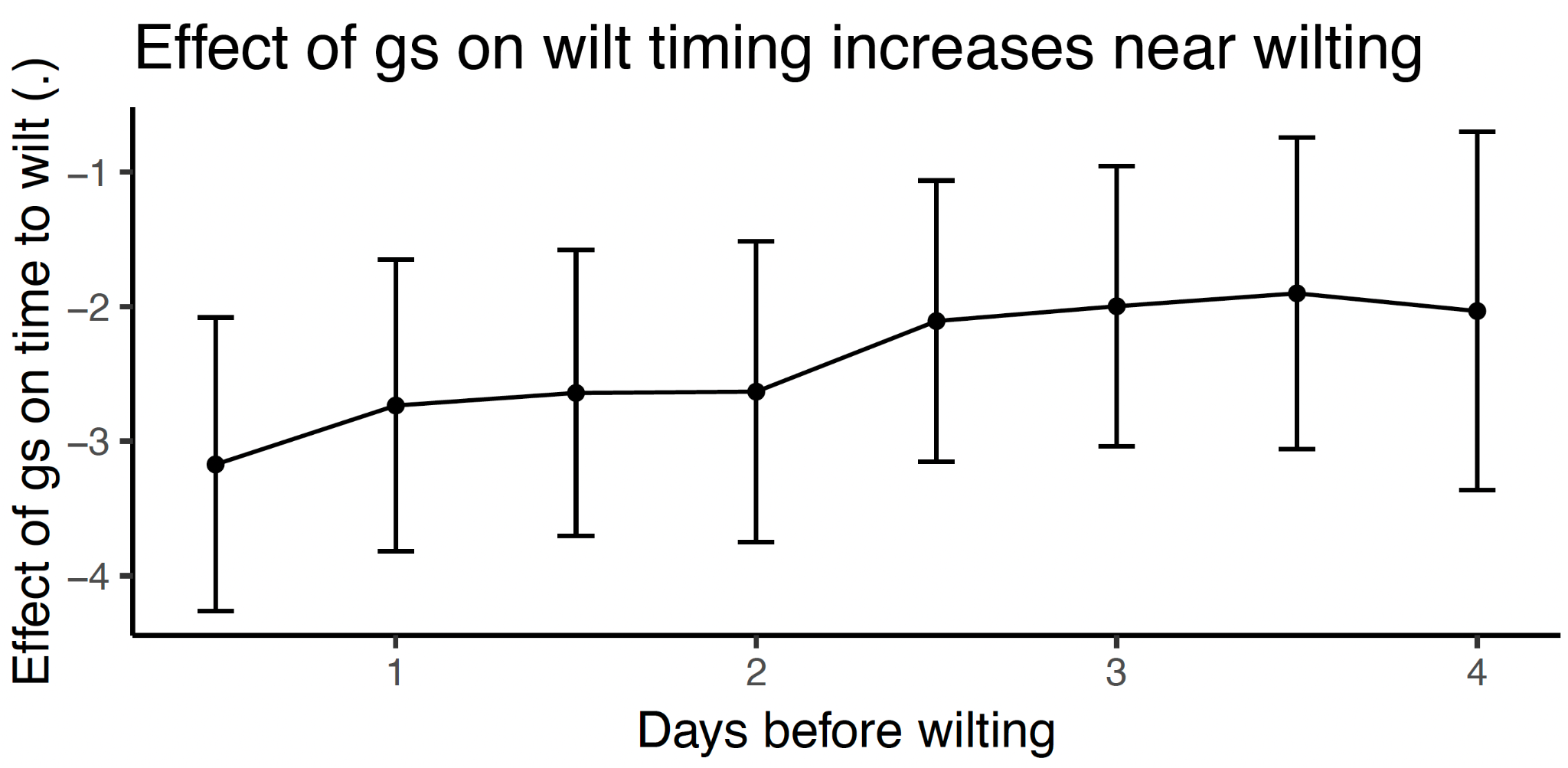

**Figure S8.** The strength of effect that stomatal conductance has prior to wilt. Stomatal conductance is more important right before wilting than it is for days prior to wilt. This suggests that stomata are not responding until significant change in water potential is beginning to be experienced by the plant.

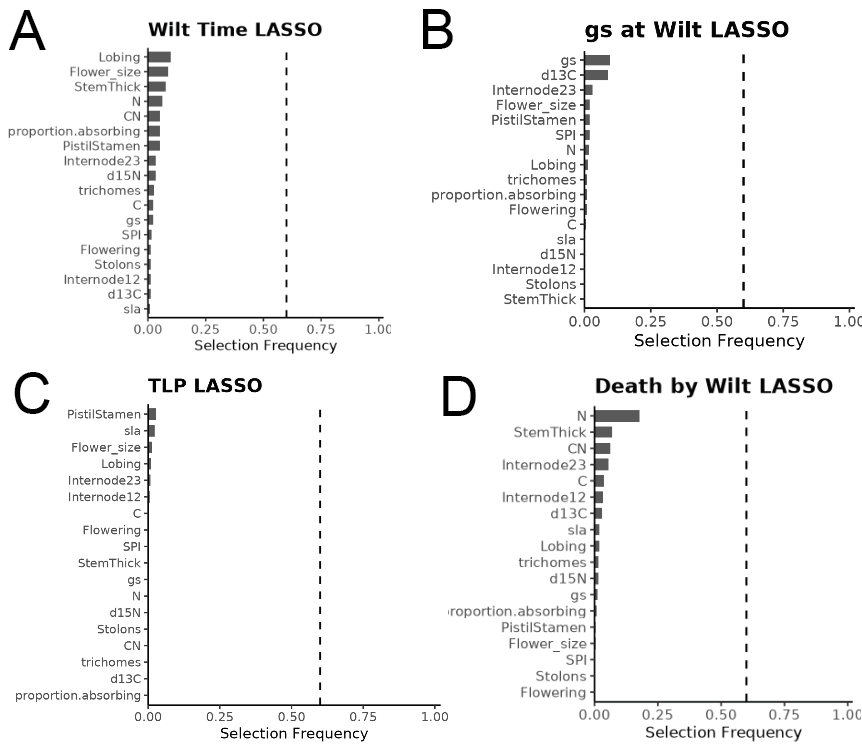

**Figure S9.** LASSO results for 18 phenotypic traits as they associate with the four drought responses. A) Wilting time/wilt resistance. B) Stomatal conductance at wilting time. C) Turgor loss point. D) The length of survival post wilt/wilt tolerance.

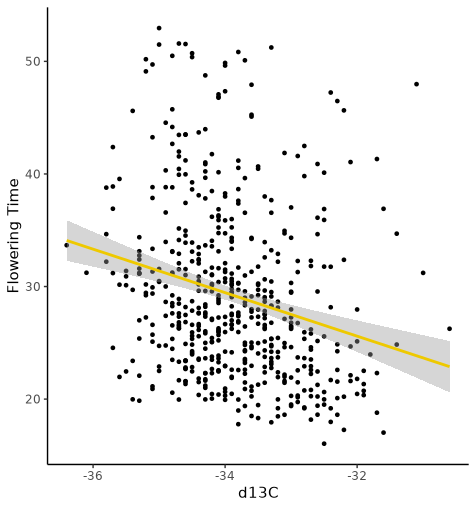

**Figure S10.** Relationship between flowering time and ^13^C (p < 0.001).
